# Discovery of an on-pathway protein folding intermediate illuminates the kinetic competition between folding and misfolding

**DOI:** 10.1101/2024.12.14.628475

**Authors:** Qing Luan, Patricia L. Clark

## Abstract

Our current understanding of protein folding is based predominantly on studies of small (<150 aa) proteins that refold reversibly from a chemically denatured state. As protein length increases, the competition between off-pathway misfolding and on-pathway folding likewise increases, creating a more complex energy landscape. Little is known about how intermediates populated during the folding of larger proteins affect navigation of this more complex landscape. Previously, we reported extremely slow folding rates for the 539 aa β-helical passenger domain of pertactin (P.69T), including conditions that favor the formation of a kinetically trapped, off-pathway partially folded state (PFS). The existence of an on-pathway intermediate for P.69T folding was speculated but its characterization remained elusive. In this work, we exploited the extremely slow kinetics of PFS unfolding to develop a double-jump “denaturant challenge” assay. With this assay, we identified a transient unfolding intermediate, PFS*, that adopts a similar structure to PFS, including C-terminal folded structure and a disordered N-terminus, yet unfolds much more quickly than PFS. Additional experiments revealed that PFS* also functions as an on-pathway intermediate for P.69T folding. Collectively, these results support a two-step, C-to-N-terminal model for P.69T folding: folding initiates in the C-terminus with the rate-limiting formation of the transient on-pathway PFS* intermediate, which sits at the junction of the kinetic competition between folding and misfolding. Notably, processive folding from C-to-N-terminus also occurs during C-to-N-terminal translocation of P.69T across the bacterial outer membrane. These results illuminate the crucial role of kinetics when navigating a complex energy landscape for protein folding.

## Introduction

Proper folding is required for protein function, yet it appears that every protein can alternatively misfold and aggregate, rather than form its native structure [1, 2]. While stably folded proteins typically adopt only one native structure (or in some cases, two [3]), the total number of possible conformations that a polypeptide chain can adopt increases exponentially with chain length [4, 5]. This higher ratio of non-native to native conformations increases the likelihood that larger proteins will populate a misfolded state [6]. Some signatures of the selective pressure to avoid stable, misfolded states are detectable in protein sequences, including selection against hydrophobic residues in large proteins [7], avoidance of long stretches of hydrophobic residues [8, 9], and, in multi-domain proteins, low sequence conservation between consecutive domains of similar structure [10]. Although it has been shown that larger, multi-domain proteins can fold co-translationally, concomitantly with vectorial appearance of the nascent chain [11], little is known regarding the folding mechanisms of larger proteins, including the competing processes that determines whether a polypeptide chain will fold to its native state or instead lead to a misfolded and/or aggregated state. This knowledge gap exists due to both the more complicated folding mechanisms of larger proteins [12] and the technical challenges of structurally characterizing subtle differences between the short-lived intermediate states that lead to folding versus misfolding.

Although less prone to misfolding in general, studies of small proteins have identified strategies that may be relevant for the successful folding of larger proteins with more complex energy landscapes. For example, even the relatively simple folding mechanisms of small proteins can include non-native interactions: Im9, an immunity protein from colicin-producing bacteria, folds by a two-state mechanism, yet its close homolog, Im7, folds by a three-state mechanism with a non-native yet on-pathway folding intermediate [13]. The difference of a few amino acid residues between Im9 and Im7 is sufficient to alter the folding mechanism without affecting the thermodynamic stability of native state, via the stabilization of an intermediate conformation with non-native, long-range contacts [14]. Similarly, a recent study of influenza nucleoprotein discovered a single point mutation that can alter the folding pathway, leading to the population of an aggregation-prone intermediate [15]. The energy landscapes of small proteins can also include unusually large energy barriers, leading to kinetic traps. A particularly extreme example of kinetic trapping is kinetic stability, where the unfolded state of a protein is more stable than its folded structure. A classic example of kinetic stability is α-lytic protease, which possesses a sufficiently high energy barrier for unfolding as to enable a native state lifetime of >1 year, despite having a native structure that is less stable than its unfolded state [16]. “Bridge” interaction formed between the N- and C-terminal regions of α-lytic protease contributes to the unusually high energy of its transition state [17]. Notably, another kinetically stable protease, SbtE, has a close homolog, ISP1, that is instead thermodynamically stable, suggesting two distinct strategies have evolved to maintain the native, functional protease structure [18]. Currently, it remains unclear how non-native interactions and/or kinetic trapped states constrain the folding of large proteins to favor the formation of productive (on-pathway) intermediate and/or avoid unproductive misfolded (off-pathway) conformations, and in particular what mechanisms enable large proteins to avoid kinetic traps that lead to misfolding.

Previously, we determined that the passenger domain of *Bordetella pertussis* pertactin (P.69T) can refold reversibly upon dilution from denaturant *in vitro* [19], despite being an unusually large protein (539 aa). Native P.69T consists of a single structural domain, a 16-rung right-handed parallel β-helix [20, 21]. We previously showed that despite a continuous hydrogen bond network throughout the β-helix, native P.69T is comprised of two segments with distinct stabilities and folding behaviors (**Figure 1A**): The N-terminus (Nt; residues 1-334) is significantly less stable than the C-terminus (Ct; residues 335-539), which enables P.69T to adopt a partially folded state (PFS) at equilibrium in intermediate concentrations of denaturant (e.g., 1-2 M guanidine hydrochloride, GdmCl) [19]. However, although populated at equilibrium, PFS is not populated on the dominant kinetic pathway between U and N, as the rates for both folding of PFS→N and unfolding of PFS→U are orders of magnitude slower than U→N folding and N→U unfolding, respectively [22]. Hence, PFS represents a misfolded state: a kinetically trapped, off-pathway conformation of P.69T (**Figure 1A**).

**Figure 1.**
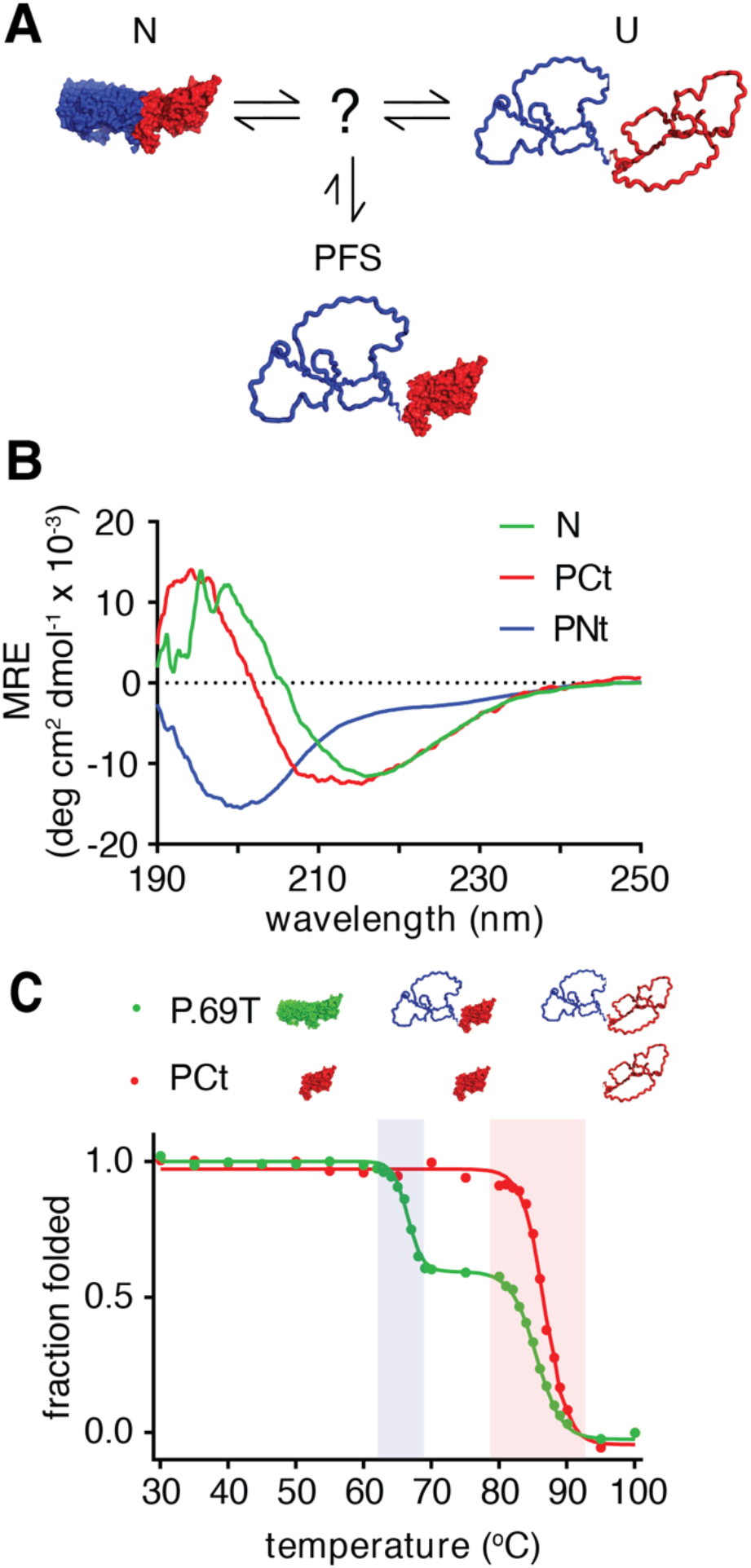
PCt, the isolated C-terminus of P.69T, closely resembles P.69T-Ct. **(A)** Schematic of P.69T folding and misfolding. PFS is off-pathway and has a much slower folding rate than full unfolded P.69T (U) and a much slower unfolding rate than native P.69T (N). An on-pathway intermediate was previously predicted [19], but not detected. The structure of P.69T (PDB ID: 1dab) [20] is shown in *blue* (Nt; residues 1-333) and *red* (Ct; residues 334-539). **(B)** Far-UV circular dichroism (CD) spectra of P.69T (*green*), PCt (*red*) and PNt (*blue*). **(C)** Thermal denaturation of P.69T (*green*) and PCt (*red*) monitored by far-UV CD spectroscopy. Data were converted to fraction of the native state CD signal at 218 nm and fit to a sigmoidal model (see **Methods**). Unfolding transitions for Nt and Ct segments are highlighted by *blue* and *red shading*, respectively.

In a past study, we added an exogenous fluorophore to either Nt or Ct to compare the folding rates for each segment of P.69T [22]. Those experiments returned the same folding rates for both segments. However, we have since discovered that the isolated N-terminus (PNt) is disordered [9, 23-25], suggesting that Nt and Ct are not equal partners in P.69T folding. In addition, an unresolved conundrum surrounding P.69T folding is the incompatibility between measured *in vitro* folding rates and relevant cellular timescales. Specifically, the fastest rate constant measured for P.69T folding *in vitro* (*k* = 5 x 10^-4^ s^-1^; τ = 30 min [22]) is significantly slower than the time needed for bacterial replication (∼20 min), which we use as a conservative upper bound for the *in vivo* folding rate [26]. Notably, P.69T folding *in vitro* is much slower than the folding of many other proteins, including other large, processively-wound (“solenoid”) proteins; e.g., the ankyrin domain of Notch [27, 28] and other proteins with β-helical structure [29, 30]. For these reasons, we have long suspected that pertactin refolding *in vitro* may be slowed by the formation of transient off-pathway folding intermediates, which the cellular environment may help avoid, thereby accelerating the P.69T folding rate.

Here we characterize the folding behavior of PCt, the isolated C-terminal segment of the pertactin passenger, and capitalize upon these results to discover an on-pathway intermediate populated during both the folding and unfolding of P.69T. This intermediate, which we call PFS*, is spectroscopically indistinguishable from PFS but thermodynamically less stable. Both PFS and PFS* consist of an unfolded, disordered Nt segment and a folded Ct. However, PFS* folds an order of magnitude faster to the native structure than to PFS, favoring the folding outcome in the kinetic competition between folding and misfolding. Strikingly, the *in vitro* P.69T folding pathway retains the C-to N-terminal folding mechanism used by the pertactin passenger domain upon its secretion across the outer membrane of *B. pertussis* [22]. Collectively, these results advance our understanding of the interplay between folding and misfolding for large proteins with complex energy landscapes.

## Results

### P.69T C-terminus (PCt) is stably folded, in contrast to PNt

For clarity, we refer to the N- and C-terminal segments within full-length P.69T as Nt and Ct, respectively, and to the isolated N- and C-terminal fragments as PNt and PCt, respectively. Previously, we showed that PNt adopts a highly expanded, disordered conformation [9, 23-25], suggesting that in P.69T the stability of Nt is dependent on Ct. Based solely on this discovery, it is tempting to speculate that P.69T folds via a simple two-step model, with an on-pathway intermediate with a disordered Nt and a natively-folded Ct, with Ct providing a template upon which Nt folds. However, while our past work identified that PFS has a stably folded Ct that is populated at equilibrium at intermediate concentrations of denaturant (1-2 M GdmCl), the rates of PFS→N folding and PFS→U unfolding are orders of magnitude slower than complete refolding (U→N) or unfolding (N→U) of native P.69T [22], demonstrating that PFS is off-pathway. One resolution of this apparent paradox would be if Ct adopts two distinct (albeit structurally similar) intermediate states, one that is on-pathway and another (PFS) that is off-pathway, with different positions on the energy landscape for P.69T folding. A crucial hypothesis raised by this model is the existence of a previously undetected on-pathway intermediate with a Ct conformation that can rapidly fold/unfold, rather than a PFS-like slow folding/unfolding state (**Figure 1A**).

As an initial approach to testing the hypothesis above, we first tested whether Ct is stably folded, independent of Nt. We cloned, expressed and purified PCt (see Methods) and found that, in stark contrast to the disorder of PNt, PCt adopts a stable, β-sheet-rich structure by far-UV circular dichroism (CD) spectroscopy (**Figure 1B**). Moreover, thermal denaturation of PCt resulted in a cooperative two-state transition with a melting temperature (*T*_m_) that closely matches the *T*_m_ for the unfolding of the Ct segment of P.69T (**Figure 1C**; *T*_m_ = 86.8°C and 85.5°C, respectively). These results demonstrate that, in direct contrast to the highly expanded, disordered structure of PNt [9, 23-25], PCt is stably folded.

As an orthogonal approach to assess the foldability of PCt, we turned to protease digestion. We have previously shown that native P.69T is largely resistant to degradation by proteinase K, except for one cleavage site within a loop that yields two fragments (apparent molecular weights of 27 and 28 kDa), which together constitute full length P.69T [19]. In contrast, under the same conditions PFS is rapidly degraded to one shorter fragment of 21 kDa, consisting of Ct, demonstrating that in the PFS Ct is stably-folded and resistant to proteinase K, whereas Nt is unfolded and rapidly degraded [19]. When PCt was subjected to proteinase K digestion under the same conditions used to digest native P.69T and PFS, PCt showed no evidence of degradation (**Figure S1**), consistent with the conclusion that PCt is capable of folding.

### PCt unfolds much more slowly than P.69T

We next tested whether PCt unfolds and refolds reversibly, and if so, whether PCt folds/unfolds rapidly (like native P.69T) or slowly (like PFS). We showed previously that it takes approximately two weeks for P.69T unfolding/refolding titrations to reach equilibrium, particularly at intermediate concentrations of denaturant where PFS is populated [19]. PCt folded status was adjusted by addition or dilution of GdmCl and monitored by tryptophan fluorescence emission spectroscopy as a function of time (**Figure 2**). PCt refolding was complete after 24 h; however, in stark contrast to native P.69T, it took longer than 6 months for the PCt unfolding titration to converge with the refolding titration, akin to the time necessary to unfold PFS [22]. Although not explored in detail here, the denaturant-dependent PCt unfolding rates observed in **Figure 2B** indicate a complex relationship between denaturant concentration and PCt unfolding kinetics. Ultimately, however, a standard two-state equilibrium between PCt folding and unfolding was established, consistent with the thermal denaturation results (**Figure 1C**). The midpoint of the PCt unfolding transition (*c*_m_, = 2.03 M) is indistinguishable from the unfolding midpoint for P.69T-Ct (*c*_m_ = 1.99 M) [19]. The *m-*values for the PCt and P.69T-Ct equilibrium unfolding transitions are likewise indistinguishable (**Figure S3**), indicating that these folding transitions are similarly cooperative. Together, these results showed that PCt adopts a conformation that is thermodynamically comparable to the Ct segment of native P.69T, but with a starkly different kinetic pathway for unfolding, more similar to the slow unfolding state as the Ct in PFS than the fast unfolding of the Ct region of native P.69T when unfolding from N→U. PCt and PFS both require >1 month to unfold in 4 M GdmCl, whereas native P.69T unfolds completely in 10 sec (**Figure 3** and [22]). These results demonstrate that, in contrast to native P.69T, both PCt and PFS adopt a kinetically trapped conformation. Related to the paradox introduced above, the much slower unfolding rate of PCt relative to P.69T-Ct excludes the possibility of an on-pathway intermediate with a PCt-like Ct.

**Figure 2.**
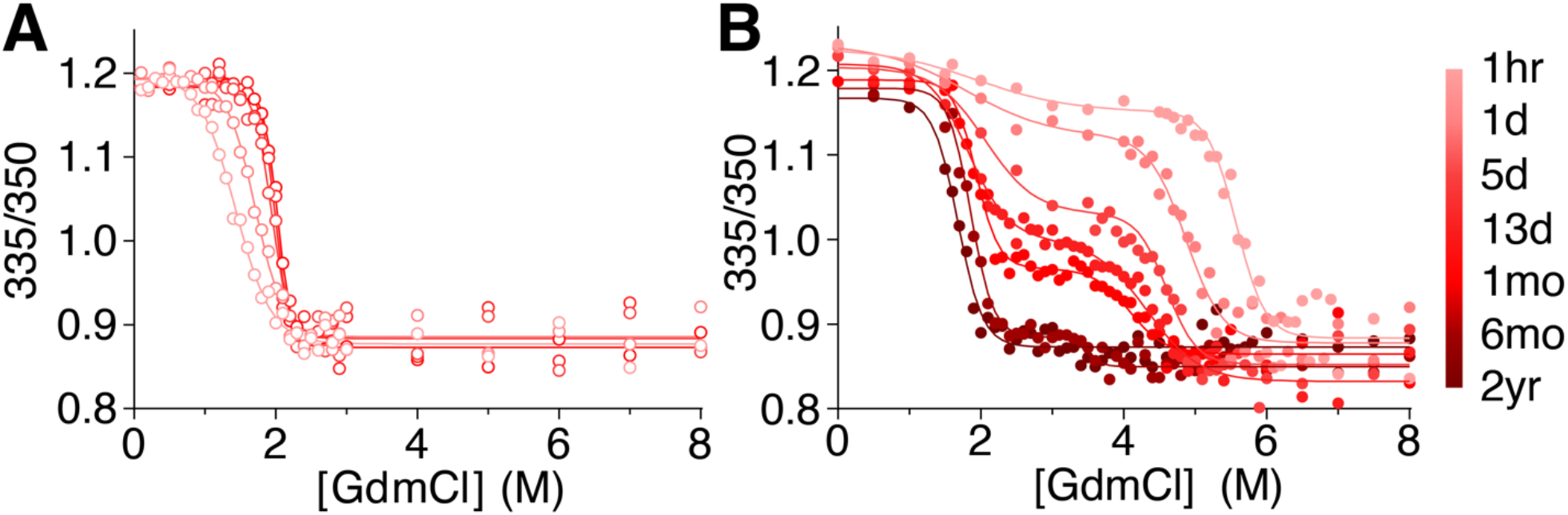
The slow approach to equilibrium during folding (**A**) and unfolding (**B**) of PCt. The equilibrium midpoint (*c*_m_) for PCt unfolding and refolding is 2.0 M GdmCl, but equilibrium is established extremely slowly. PCt was refolded from 8 M GdmCl (**A**, *open symbols*), or unfolded (**B**, *filled symbols*) to the indicated final GdmCl concentration, measured as the ratio of tryptophan emission intensities at 335 and 350 nm. Lines represent the fit to single or double sigmoidal models (see **Methods**).

**Figure 3.**
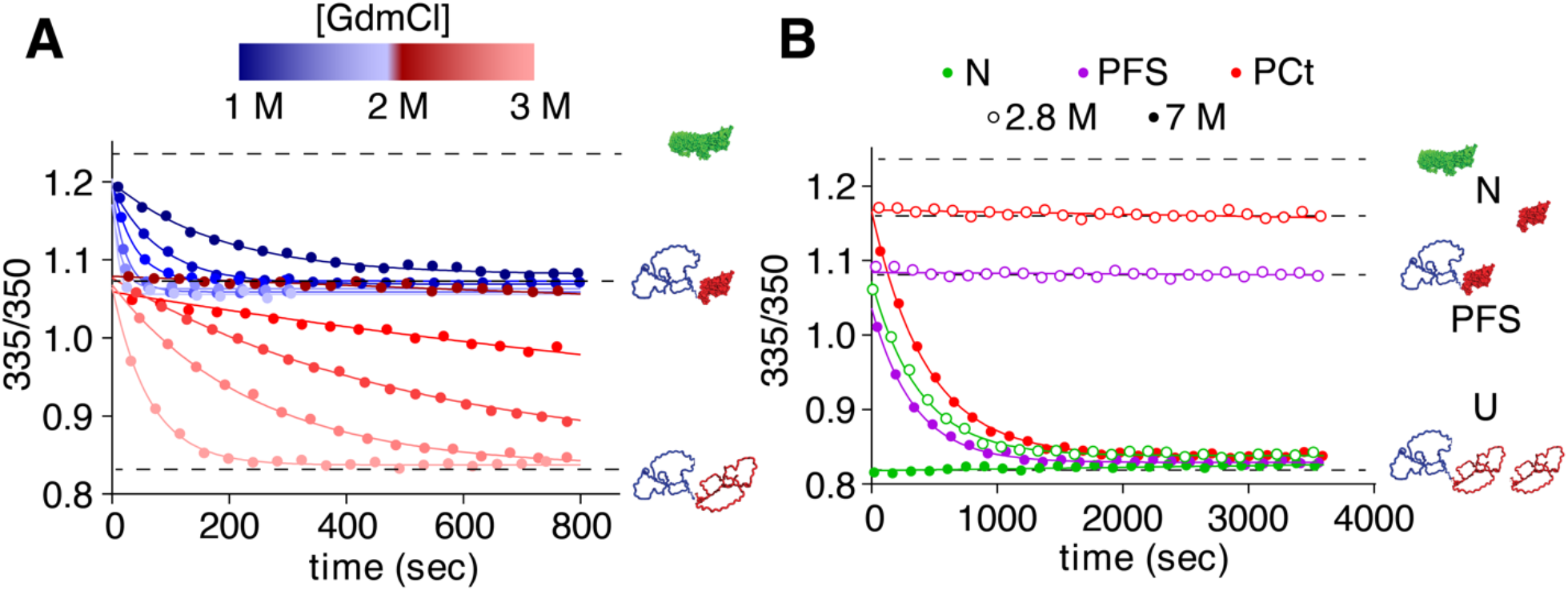
For native P.69T, unfolding of Nt accelerates the unfolding of Ct. **(A)** Unfolding kinetics of P.69T as a function of GdmCl concentration. Each data set was fit to either a single-or double-exponential equation (see **Results** and **Methods**). Rate constants for unfolding are summarized in **Table S1.** **(B)** Unfolding of native P.69T (*green*), PFS (*purple*) and PCt (*red*) at two GdmCl concentrations (*open circles*: 2.8 M; *filled circles*: 7 M), showing that native P.69T unfolds orders of magnitude more rapidly than PFS and PCt.

### Unfolding of Nt accelerates Ct unfolding

The slow PCt unfolding rate described above suggests that the presence of Nt and/or its unfolding accelerates the unfolding of the Ct region within native P.69T. To test this, we first used changes in tryptophan fluorescence emission to measure the denaturant-dependent unfolding rates of Nt and Ct within native P.69T, exploiting the distinct stabilities of Nt and Ct to make these measurements. Previously, we reported overall unfolding rate constants for native P.69T and PFS [22], but the unfolding rates for the individual Nt and Ct segments were not characterized. Over a range of [GdmCl] from 1.1-2.1 M, the Nt region of native P.69T unfolds, leading to the PFS conformation (**Figure 3A**) [19]. Nt unfolding kinetics fit well to a single exponential model (**Figure 3A, Table S1**). As expected, as [GdmCl] increased, the rate constant for Nt unfolding increased, eventually occurring entirely within the dead time of our experimental setup (10 s). Nt unfolding corresponded to ∼40% of the total fluorescence change upon unfolding, a percentage consistent with the number of tryptophan residues in each segment (Nt = 3/7 Trp residues; Ct = 4/7 Trp residues). Below 2.1 M GdmCl, Ct remained folded. Above 2.3 M GdmCl, both Nt and Ct unfolded. At these higher denaturant concentrations, the unfolding kinetics fit best to a double exponential model (**Figure 3A, Table S1**). The fast phase and slow phase consist of approximately 40 and 60%, respectively, of the total change in fluorescence upon unfolding. The fast phase was completed within the dead time of the experimental setup (10 s), consistent with the Nt unfolding rate constant extrapolated from measurements at lower GdmCl concentrations. The Ct unfolding rate constant increased with increasing GdmCl concentration, and above 3.5 M GdmCl became too fast to accurately quantify using our experimental setup. These results are consistent with a model where the unfolding rate of Nt is faster than Ct under all conditions tested.

To determine whether the presence of Nt affects Ct unfolding, despite Nt unfolding more rapidly than Ct, we directly compared the rate constants for P.69T and PCt unfolding. Because the rates of these unfolding processes differ by orders of magnitude, it is difficult to measure the unfolding of both native P.69T and PCt under identical conditions. To address this, we selected two denaturant concentrations, 2.8 M and 7 M GdmCl, as representative scenarios. As discussed above, in 2.8 M GdmCl native P.69T unfolding was best fit to a double exponential model, with an unfolding rate constant for native P.69T-Ct of 2.61×10^-3^ s^-1^ (**Figure 3B**). In contrast, no detectable unfolding of PCt occurred within one hour at 2.8 M GdmCl (**Figure 3B, Table S2**). In 7 M GdmCl, native P.69T unfolded to U entirely within the dead time (<10 s), whereas PCt unfolding kinetics fit well to a single exponential with *k* = 2.29×10^-3^ s^-1^ (**Figure 3B, Table S2**). Hence, native P.69T unfolds faster than PCt over a wide range of conditions, indicating that Nt accelerates the unfolding of the Ct segment of native P.69T.

To test whether the conformation of Nt – or just its presence, regardless of conformation – affects Ct unfolding, we compared the N→U unfolding rate constants measured above to the rate constant for PFS unfolding under identical conditions. To form PFS, native P.69T was incubated in 1.5 M GdmCl overnight. Subsequent unfolding of PFS in 2.8 M or 7 M GdmCl confirmed that the PFS unfolding rate constant is indistinguishable from the rate constant for PCt unfolding (**Figure 3B, Table S2**), indicating that the presence of an unfolded Nt (as in PFS) is insufficient to accelerate the unfolding rate of Ct in P.69T. In contrast, unfolding of the native fold of Nt led to a 100-1,000-fold acceleration of Ct unfolding under the conditions tested here.

### PFS* is an on-pathway unfolding intermediate

As shown above, the Nt segment of P.69T unfolds faster than the Ct segment, which suggests that P.69T unfolding from N to U may include the transient formation of a PFS-like intermediate. However, as described above, unfolding of PFS to U is orders of magnitude slower than unfolding of native P.69T to U (**Figure 3B, Table S2**) [22], excluding PFS as the on-pathway unfolding intermediate between N and U. To resolve this paradox, we hypothesized that unfolding of the folded Nt segment leads to a transient structure that resembles PFS spectroscopically but is not yet kinetically trapped. To test this hypothesis, we designed a double-jump “denaturant challenge” experiment (**Figure 4A**), taking advantage of the extremely slow unfolding rate of the kinetically trapped PFS conformation. Native P.69T was first incubated in 1.5 M GdmCl, which is sufficient to rapidly unfold Nt (**Figure 3A**). After 3 min, additional GdmCl was added, to 4.75 M. If the 3 min incubation in 1.5 M GdmCl was sufficient to not only unfold Nt but also form PFS, the subsequent transfer to 4.75 M GdmCl should lead to no observable unfolding to U within a few hours [22]. Surprisingly, however, the brief incubation of P.69T in 1.5 M GdmCl led to rapid unfolding to U within the dead time of transfer to 4.75 M GdmCl (**Figure 4B**). These results reveal that, upon unfolding of Nt, P.69T initially adopts a denaturant-sensitive, transient conformation that is spectroscopically indistinguishable from PFS, which we term PFS*, prior to forming PFS, which is kinetically trapped.

**Figure 4.**
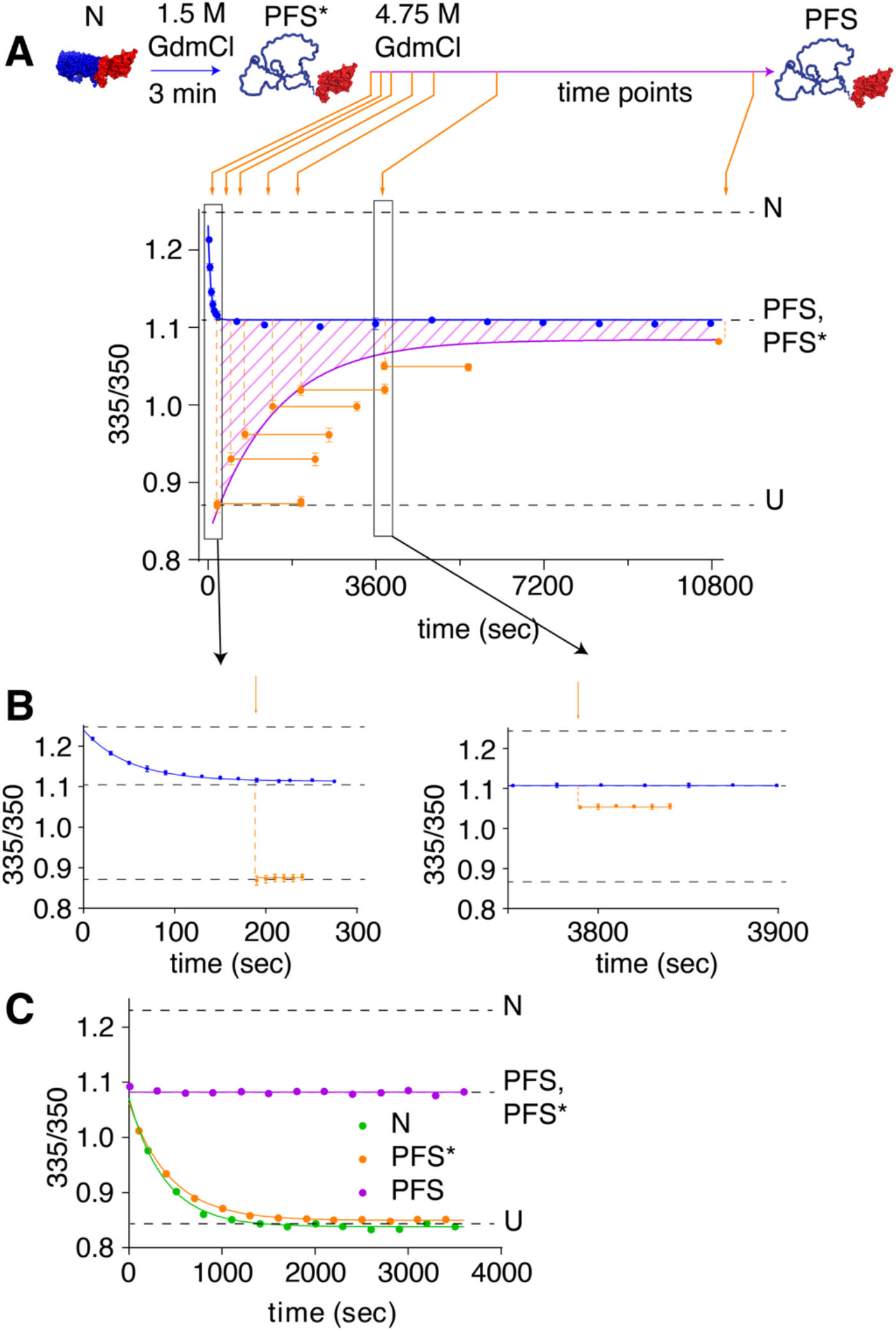
Discovery of PFS*, an on-pathway unfolding intermediate. **(A)** Kinetics of PFS formation in the denaturant challenge experiment, monitored by tryptophan fluorescence emission spectroscopy. Incubating native P.69T in 1.5 M GdmCl for 3 minutes led to unfolding of Nt within 300 s (blue line; see also panel **B** and **Figure 3A**). At various tim points after the initiation of unfolding (indicated by orange arrows), the impact of the addition of 4.75 M GdmCl on P.69T unfolding was evaluated. The fraction of P.69T molecules in the PFS conformation, which unfolds extremely slowly in 4.75 M GdmCl (orange data points), increased as a function of time and fit to a single exponential model (purple line). The hatched region between the blue and purple curves represents the fraction of P.69T molecules in an alternative conformation, PFS*, that resembles PFS but is more susceptible to unfolding. The population of molecules in the PFS* conformation decreased as a function of incubation time in 1.5 M GdmCl. Error bars represent the standard deviation of three independent experiments; in several instances, the error bar is smaller than the height of the data point. **(B)** Examples of denaturant challenge experiments at 3 min and 63 min after the beginning of Nt unfolding. **(C)**PFS* unfolds in 2.8 M GdmCl at a rate similar to native P.69T, but much faster than PFS.

To track the fate of PFS* – specifically, its rate of conversion to the kinetically trapped PFS – we incubated native P.69T in 1.5 M GdmCl for different amounts of time prior to the 4.75 M GdmCl denaturant challenge described above. We found that P.69T started to exhibit resistance to unfolding in 4.75 M GdmCl as soon as 8 min after addition of 1.5 M GdmCl. The fraction of P.69T resistant to rapid unfolding in 4.75 M GdmCl increased as a function of time (**Figures 4A & S4**) and fit well to a single exponential model with a rate constant of 7.1×10^-4^ ± 6×10^-5^ s^-1^. This result indicates that conversion of PFS* to PFS is spontaneous in 1.5 M GdmCl and represents the rate limiting step of converting native P.69T to PFS, which is kinetically trapped.

The results above demonstrate that in 1.5 M GdmCl, native P.69T initially unfolds to PFS*, prior to forming PFS. These results do not, however, discern whether the PFS* intermediate is populated during the transition from N and U, or – like PFS – is an off-pathway misfolded state. To test whether PFS* is populated during P.69T unfolding, we compared the unfolding kinetics of native P.69T, PFS and PFS* in 2.8 M GdmCl. P.69T was incubated in 1.5 M GdmCl for 3 minutes or overnight, to form PFS* or PFS, respectively. In 2.8 M GdmCl, the unfolding of P.69T from N overlapped with that of PFS*, whereas there was no detectable unfolding of PFS over 1 h (**Figure 4C, Table S2**). A similar experiment monitored by far-UV CD spectroscopy yielded comparable results (**Figure S5**). These results demonstrate that formation of PFS* is compatible with the unfolding of P.69T from N to U. We propose that native P.69T unfolding proceeds in a stepwise fashion, initially by unfolding Nt to PFS*, followed by unfolded to U. In this manner, PFS* serves as an on-pathway unfolding intermediate that, if populated for sufficient time (e.g., as in 1.5 M GdmCl), slowly converts to the kinetically trapped PFS (**Figure 6**).

### PFS* likely also represents the on-pathway intermediate for P.69T folding

The discovery that PFS* is an on-pathway intermediate during P.69T unfolding raises the question of whether PFS* is also populated during P.69T folding from U. As a first test of whether a PFS*-like state lies between U and N, we determined the folding rate of PFS*→N. We previously showed that PFS→N folding is orders of magnitudes slower than U→N folding [22], and to date, we have not identified a P.69T intermediate that folds to the native P.69T structure faster than U. Building upon the double-jump denaturant challenge experiment described above, we incubated native P.69T in 1.5 M GdmCl for just 3 minutes, to form PFS*, before jumping back to 0.5 M GdmCl to observe refolding to N. Consistent with the model that PFS* is an on-pathway intermediate for U→N folding, refolding from PFS*→N led to the fastest rate ever observed for refolding to native P.69T (5.3×10^-3^ ± 4×10^-4^ s^-1^, **Figure 5A,B, Table S3**), 10-fold faster than refolding from U→N (5.8×10^-4^ ± 3×10^-5^ s^-1^, **Figure 5A,B, Table S3**) and 100-fold faster than PFS→N [22]. The rate acceleration observed for starting from PFS* as opposed to U (or PFS) supports a model where the PFS* conformation is an on-pathway intermediate between U and N.

**Figure 5.**
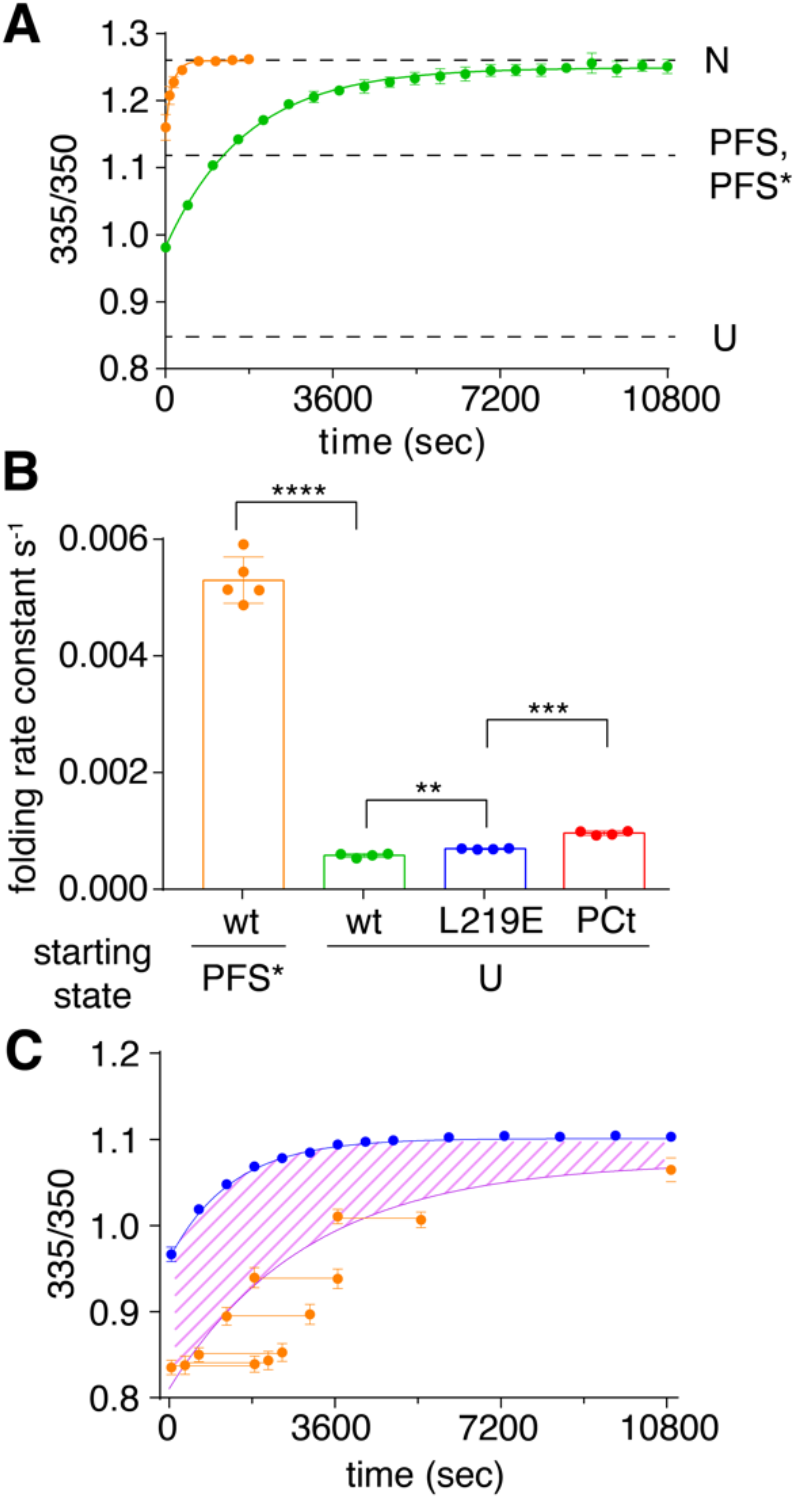
PFS* is an on-pathway folding intermediate for P.69T folding from U→N. **(A)** Faster kinetics for folding to the P.69T native structure when starting from PFS* (*orange*; *k*_f_ = 5.3×10^-3^ ± 4×10^-4^ s^-1^ SEM) versus U (*green*; *k*_f_ = 5.8×10^-4^ ± 3×10^-5^ s^-1^ SEM). **(B)** Comparison of folding rate constants in 0.5 M GdmCl for PFS* (*orange*), fully unfolded P.69T (*green*), P.69T-L219E (*blue*) (see **Fig S8A**), and PCt (*red*) (see **Fig S8B**). Error bars represent the standard deviation of independent experiments. Student’s t-test: **, P <0.01; ***, P<0.001; **** P <0.0001. **(C)** Kinetics of PFS* formation during refolding of P.69T-L219E. P.69T-L219E was fully unfolded in 7 M GdmCl. Refolding was initiated by dilution into 0.5 M GdmCl; this corresponds to *t* = 0 s. The overall progress of refolding, as measured by the 335/350 nm fluorescence emission ratio, is shown in *blue*. Alternatively, at each indicated time point, an aliquot of the refolding sample was transferred to 4.25 M GdmCl. *Orange points* indicate the impact of this “denaturant challenge” on P.69T-L219E fluorescence ratio at different refolding timepoints. The fraction of P.69T-L219E molecules in the PFS conformation, which unfolds extremely slowly in 4.25 M GdmCl, increased as a function of time and fit well to a single exponential model (*purple line*). The *hatched region* between the blue and purple curves represents the fraction of P.69T-L219E molecules in the PFS* conformation (see **Fig S8**). Error bars represent SEM of three independent experiments. The height of some error bars is smaller than the size of the datapoints.

**Figure 6.**
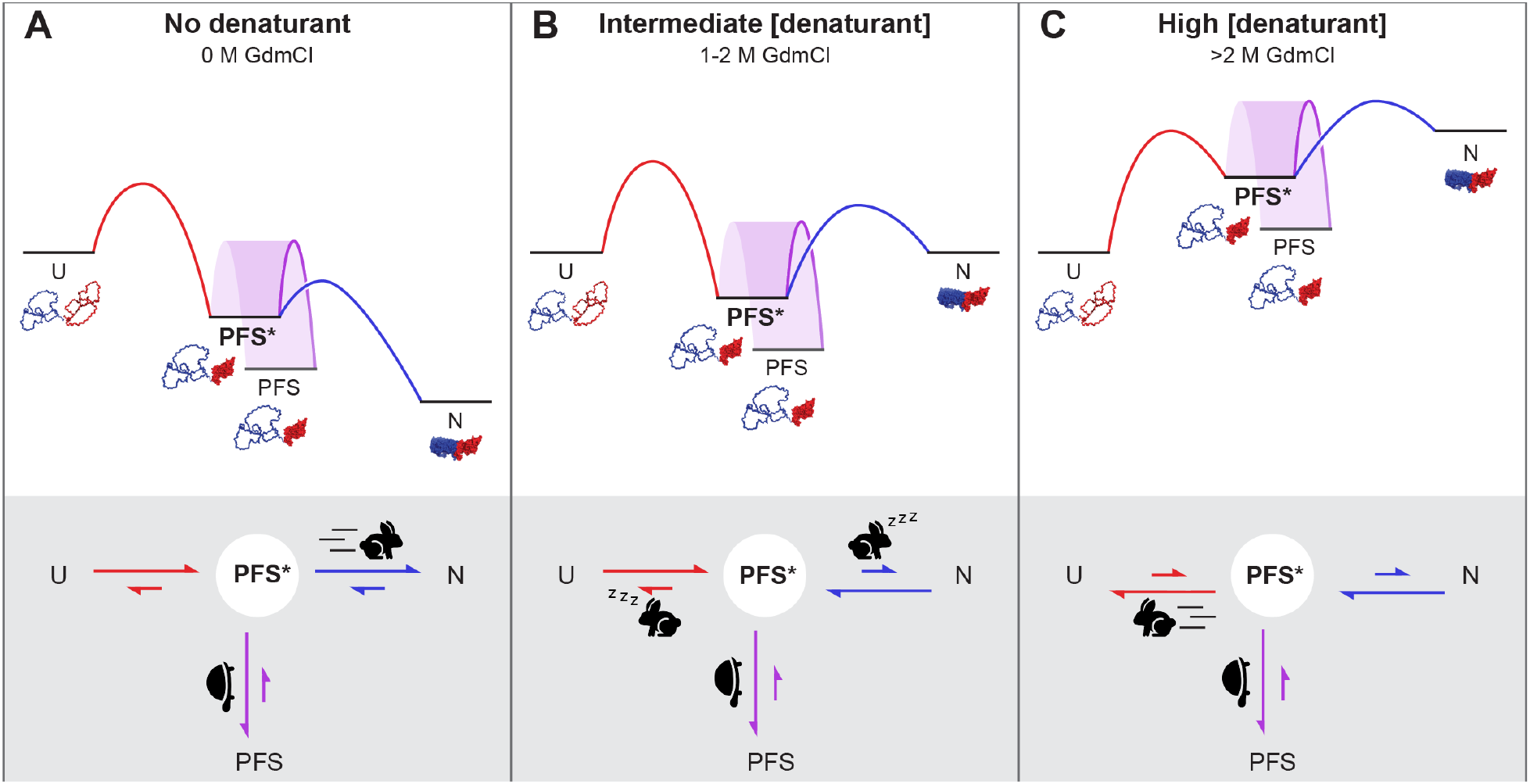
Schematic illustrating features of the energy landscape and kinetic pathway for P.69T folding and unfolding at different denaturant concentrations, with an analogy to Aesop’s fable *The Tortoise and the Hare*. An on-pathway intermediate state, PFS*, lies between the native (N) and fully unfolded (U) states. PFS* spontaneously converts to the more stable, off-pathway misfolded state, PFS, at a rate that is largely insensitive to denaturant. Note that the energy of the unfolded state has been normalized to facilitate comparisons across the three panels. **(A)** Under native or near-native conditions, folding is favored. The Ct segment of P.69T first folds slowly to the on-pathway intermediate PFS*, after which rapid folding of Nt (illustrated as a running hare) kinetically outcompetes the conversion of PFS* to PFS (illustrated as a tortoise), leading to formation of the native P.69T β-helix structure. **(B)** In contrast, at intermediate denaturation conditions PFS* is more stable than either N or U, but PFS is even more stable. Moreover, while the energy barrier between PFS* and PFS remains steady (as a tortoise), the energy barriers that separate PFS* from either N or U are more sensitive to denaturation and now higher than the energy barrier to PFS (illustrated as sleeping hares), and thus the off-pathway misfolded state, PFS, predominates. **(C)** At high denaturation conditions (>2 M GdmCl), unfolding is favored. Native P.69T-Nt first unfolds to form PFS*, followed by rapid unfolding of P.69T-Ct (illustrated as a running hare), which occurs faster than the conversion of PFS* to PFS (illustrated as a tortoise), enabling unfolding to U to outcompete misfolding.

Because the folding of PFS*→N is 10-fold faster than U→N folding, PFS* does not accumulate during refolding and is therefore difficult to study directly during refolding from U. As an alternative approach, we examined whether a PFS*-like structure is also populated during the folding of P.69T bearing a point mutation, L219E, that introduces a charged residue at a buried site in the Nt segment of native P.69T. Due to this mutation, P.69T-L219E adopts a PFS-like structure even under native conditions (**Figures S1 & S6**). We subjected this mutant construct to a double-jump denaturant challenge as described above: Denatured P.69T-L219E was first diluted to 0.5 M GdmCl to start refolding. At different time points, the refolding reaction was challenged with >4 M GdmCl. We found that that the fraction of P.69T-L219E prone to rapid unfolding upon transfer to >4 M GdmCl was initially high but decreased gradually over time (**Figures 5C & S7A**), consistent with the initial formation of the PFS* intermediate during P.69T-L219E refolding, followed by its slow conversion to PFS. Notably, the rate constant for conversion of PFS*→PFS is largely insensitive to GdmCl concentration (3.3×10^-4^ ± 3×10^-5^ s^-1^ for P.69T-L219E in 0.5 M GdmCl and 7.1×10^-4^ ± 6×10^-5^ s^-1^ for wild type P.69T in 1.5 M GdmCl) (**Figure 4A**). To slow down the PFS*→U unfolding rate to a measurable regime, we repeated the experiment but challenged with 2.8 M GdmCl, instead of >4 M (**Figure S7B**,**C, Table S2**). Again, the unfolding rate constant for P.69T-L219E in the PFS* conformation was indistinguishable from wild type PFS* (**Figure S7C**), supporting a model where a PFS*-like conformation is populated during U→N refolding of both P.69T-L219E and wild type P.69T.

Based on the findings above, we hypothesized that P.69T folds in a two-step process from C-to-N-terminus (**Figure 6A**). Ct folds first, forming the intermediate PFS*, which catalyzes the folding of Nt. Under this model, the first step (U→PFS*) is rate limiting; thus, the rate constant from U→N should be comparable to the rate constant from U→PFS*, and both rate constants should be at least 10-fold slower than folding of Nt from PFS*→N. To test this hypothesis, we measured the U→N folding rates of P.69T-L219E and P.69T. Each protein was fully unfolded in 7 M GdmCl before diluting to 0.5 M GdmCl to initiate folding. Because PFS* is spectroscopically indistinguishable from native P.69T-L219E, the folding kinetics monitored by tryptophan fluorescence reflect the folding rates from fully unfolded state to PFS*. We found that in all cases, the folding kinetics fit well to a double exponential model, with a fast phase completed in the dead time (10 s). We used the rate constant of the slow phase to represent the rate limiting step of folding. We found that the P.69T-L219E folding rate constant was 7.2×10^-4^ ± 1×10^-5^ s^-1^, similar to the P.69T folding rate constant (5.8×10^-4^ ± 3×10^-5^ s^-1^) (**Figures 5B & S8A, Table S3**). Both of these folding rate constants are an order of magnitude slower than the folding rate from PFS*→N. Collectively, these observations support the two-step C-to-N-terminus folding model described above. Notably, the rate difference between U→N folding of P.69T and P.69T-L219E was small, but statistically significant, suggesting that a foldable Nt slows down the folding of Ct, as compared to a construct with an Nt that is incapable of folding. To further elucidate the impact of Nt on the rate of Ct folding, we measured the folding rate of PCt U→N (**Figure S8B**) and found that PCt folding (9.5×10^-4^ ± 4×10^-5^ s^-1^) was significantly faster than both P.69T and P.69T-L219E (**Figure 5B, Table S3**), suggesting the both the presence and the foldability of Nt retards folding of the covalently linked Ct.

## Discussion

Protein folding is a cooperative process, requiring coordinated interactions between hundreds of atoms. The rapidity and cooperativity of protein folding makes it challenging to analyze folding intermediates, a challenge that is exacerbated when folding and misfolding occur simultaneously. Historically, the complications of competing, off-pathway misfolding reactions have led to a strong focus on model proteins with simple folding pathways and negligible competition from misfolding [31]. Yet this focus has led to relatively little insight into how larger proteins successfully navigate the regions of their folding energy landscapes that lie close to the intersection between folding and misfolding.

Here we show that the folding pathway of P.69T, the β-helical passenger domain of the 539 aa *B. pertussis* autotransporter protein pertactin, includes a previously uncharacterized intermediate, PFS*, that lies at the intersection between misfolding and folding to the native structure (**Figure 6A**). PFS* appears to be an obligatory intermediate for P.69T folding, where the folding rate of Nt folding outcompetes its conversion to an off-pathway kinetically trapped state, PFS, and favors folding to the P.69T native structure. Conditions that prolong the lifetime of PFS* increase the likelihood of its conversion to PFS (**Figure 6B**). PFS and PFS* both possess a folded C-terminus and disordered N-terminus and are indistinguishable using standard spectroscopic approaches, which for years has complicated the detangling of the P.69T folding and misfolding pathways [19, 22]. However, results from the double-jump experiments reported here show that upon dilution from denaturant, PFS* is formed more rapidly than PFS, and PFS* is converted to the native structure many orders of magnitude more quickly than the conversion of PFS→N.

A P.69T folding pathway that begins with the rate-limiting folding of the C-terminus to PFS* followed by rapid folding of Nt stands in contrast to a previous model for P.69T folding, drawn from site-specific fluorescent labeling measurements, that the N- and C-termini fold in a concerted mechanism [22]. Our discovery that Nt folding is dependent on Ct folding, yet orders of magnitude faster, enabled this update. Crucially, C-to-N-terminal folding of P.69T is also observed *in vivo*, during C-to-N-terminal translocation across the outer membrane (OM) of *B. pertussis* [32]. It is striking that, even without the spatial constraints of membrane translocation, P.69T folding *in vitro* retains this C-to-N-terminal folding vector. A similar result was predicted by steered molecular dynamics simulations [33]. Nevertheless, the rate constant for folding of P.69T *in vitro* reported here is still much slower than the expected folding rate *in vivo* [26], suggesting that the cellular environment further accelerates P.69T folding. Potentially, folding vectorially from C-to-N-terminus as P.69T is translocated across the OM reduces unproductive long-range interactions between Nt and Ct during folding. The observation that the presence of Nt slows down the folding of Ct (**Figure 5B**) supports this model.

Another distinctive aspect of P.69T folding *in vivo* is the lack of folding that occurs immediately after P.69T synthesis, when pertactin is translocated through the inner membrane (IM) into the periplasm [34]. During IM translocation, the first portion of P.69T to enter the periplasm is the disordered Nt, positioning it to retard the folding of Ct when it appears in the periplasm, as we show here also occurs during refolding *in vitro* (**Figure 5B**). In contrast, during translocation across the OM, P.69T-Ct appears first, which could enable Ct to fold at a much faster rate. Although P.69T is a large protein of 539 aa, its native β-helix structure is dominated by local contact order (CO; absolute CO = 17.6, relative CO = 0.0327), leading to a predicted folding rate many orders of magnitude faster than the measured *in vitro* folding rate observed [35, 36]. The folding mechanism proposed here highlights the distinct contributions of the N- and C-terminus to P.69T folding, contributions that are retained even in the absence of vectorial translocation across cell membranes. Collectively, these results demonstrate that both the innate protein folding properties observed in the test tube and the cellular context for folding contribute to efficient folding of a protein to its native structure, including the avoidance of off-pathway misfolded structures.

## Materials & Methods

### Cloning

A plasmid constructed previously [9] was used for expression of the pertactin passenger domain (P.69T) in the cytoplasm of *E. coli*. Truncations and point mutants were generated by site-directed mutagenesis and subcloning of this plasmid. The sequence of each expression plasmid was confirmed by Sanger sequencing. Amino acid residue numbers are consistent with the pertactin crystal structure (PDBID 1DAB) [20]. A335 was previously determined as the N-terminal residue of Ct [19], however, H334 was included in the PCt construct to provide a spacer between A335 and the N-terminal methionine residue introduced by the start codon.

### Protein purification

Wild type and mutant pertactin constructs were expressed in *E. coli* BL21(DE3)pLysS and proteins were purified from inclusion bodies as described previously [19]. Briefly, a single colony of transformed *E. coli* BL21(DE3)pLysS was used to inoculate an overnight culture in LB medium with 50 mg/L ampicillin (LB-Amp). One milliliter of this overnight culture was used to inoculate 100 mL LB-Amp, followed by incubation at 37ºC with shaking to OD_600_ = 1. A portion (40 mL) of this culture was then used to inoculate 2 L LB-Amp, which was incubated at 37ºC with shaking until OD_600_ = 0.6-0.8. Expression was induced by addition of 500 μM isopropyl β-D-1-thiogalactopyranoside (IPTG). Cells were incubated at 37ºC with shaking for an additional 2 h, at which point 2 mM ethylenediaminetetraacetic acid (EDTA) and 0.5 mM phenylmethylsulfonyl fluoride (PMSF) were added and cells were harvested by centrifugation at 3,750x*g*. Cell pellets were stored -80ºC. Cell pellets were thawed and resuspended in cell lysis buffer (50 mM TrisHCl pH7.5, 100 mM NaCl) with protease inhibitors and 1 mg/mL lysozyme, then lysed by sonication. The cell lysate was centrifuged at 12,000x*g* for 20 min, and the insoluble fraction from the centrifugation was washed with 1% Triton X-100 in cell lysis buffer and solubilized with 25 mL 6 M GdmCl in cell lysis buffer by shaking at 4ºC overnight. Solubilized inclusion bodies were diluted with dialysis buffer (50 mM TrisHCl pH8.0) at a 1:2 ratio and dialyzed against 10 L dialysis buffer for two days to remove denaturant. Dialysis buffer was refreshed once after one day to further lower the residual denaturant concentration to <0.03 mM. The dialysate was clarified by centrifuging at 12,000x*g* for 20 min and purified by chromatography, as described below.

P.69T and P.69T-L219E was first purified by anion exchange chromatography (Source 15Q, Cytiva), by loading onto the column in 50 mM TrisHCl pH8 and eluting with a gradient of 0 to 150 mM NaCl in 50 mM TrisHCl pH8. Native P.69T eluted at 6-8 mS/cm and P.69T-L219E eluted at 12-14 mS/cm. Ion exchange chromatography fractions enriched in pertactin constructs were pooled and purified further using size exclusion chromatography (Superdex 200, Cytiva) equilibrated with SEC buffer (50 mM TrisHCl pH7.5, 100 mM NaCl). PCt was purified using only size exclusion chromatography, in SEC buffer. Final purity of the protein (>99%) was verified by Coomassie stained SDS-PAGE. Purified proteins were concentrated using centrifugal filters (Sartorius Vivaspin Turbo 15, 10K MWCO) in storage buffer (25 mM TrisHCl pH7.5, 50 mM NaCl) followed by aliquoting, flash freezing and storage at -80ºC.

### Far-UV circular dichroism spectrometry

Far-UV CD spectra, thermal denaturation curves and kinetic traces were collected using a J-815 CD spectropolarimeter (Jasco) as previously described [23]. For each spectrum, 5 μM protein in 25 mM sodium phosphate pH7.5 buffer was measured in 1 mm quartz cuvette (Starna). Data were obtained with a bandwidth step of 1 nm and data integration time of 1 s and represented as the average of a triplicated measurement of the same sample. Thermal denaturation data and kinetic traces were monitored at 218 nm and measured in 2 mm quartz cuvette (Starna). Thermal denaturation curves were collected using 5 μM P.69T and P.69T-L219E or 10 μM PCt in 25 mM sodium phosphate pH7.5 buffer. The temperature ramp rate was 2.5 min/°C and data integration time was 4 s. Thermal denaturation curves were fit to a single or double sigmoidal model and the melting temperature (T_m_) was defined as the inflection point of the sigmoidal curve (see Data analysis and statistics, below). Kinetic traces were collected using 2.5 μM protein. Data acquisition interval was 5 s and integration time was 4 s.

### Proteolytic digestion

P.69T and variants (5 μM each) were digested with 2 μg/ml proteinase K (Sigma) for 24 h at room temperature in 50 mM TrisHCl pH8.8, 7.5 mM CaCl_2_. For PFS, P.69T was incubated in 1 M GdmCl at room temperature overnight before addition of protease supplemented with 1 M GdmCl. Proteolysis reactions were quenched by boiling. Digested fragments were resolved by Coomassie stained SDS-PAGE. Images were processed using Fiji [37].

### Fluorescence spectroscopy

All fluorescence measurements were collected using a Fluorolog-QM spectrofluorometer (Horiba). Samples were measured in a 10 mm quartz cuvette (Starna) at 20°C. Data was collected using the instrument’s Felix-GX software and corrected for excitation fluctuations and emission offset using the built-in quanta lookup table provided by the manufacturer.

### Equilibrium time course unfolding and refolding

The approach to equilibrium for PCt unfolding and refolding was monitored by changes in tryptophan fluorescence emission. For unfolding, 200 nM PCt was incubated in 0-8 M GdmCl, 25 mM TrisHCl pH7.5, 50 mM NaCl in 5 ml Protein LoBind tubes (Eppendorf). For refolding, PCt were first fully unfolded in 8 M GdmCl, 25 mM TrisHCl pH7.5, 50 mM NaCl, then diluted with 25 mM TrisHCl pH7.5, 50 mM NaCl to 4,800 μL, to a final concentration of 200 nM PCt and 0.1-8 M GdmCl in 5 ml Protein LoBind tubes (Eppendorf). To avoid time-dependent photobleaching, three strategies were employed: (i) samples were stored in the dark, (ii) at each indicated time point, the fluorescence emission of a fresh aliquot at each concentration was measured, and (iii) fluorescence emission changes were calculated as a ratio of two values, rather than relying on changes in absolute intensity.

At various time points (unfolding: 1 hr, 1 d, 5 d, 1 mo, 5 mo, 26 mo; refolding: 1 h, 4 h, 1 d, 2 d, 6 d), the tryptophan emission spectrum was collected, with excitation at 280 nm (slit width 2 nm) and emission measured from 315-380 nm (slit width 5 nm), with 0.2 s exposure at a wavelength scan step of 1 nm. The same corrected fluorescence values were collected as described above, and the ratio between fluorescence emission intensities at 335 nm and 350 nm (referred to as 335/350) was calculated at each time point for each GdmCl concentration. The resulting titration curves were fit to a single or double sigmoidal model and the denaturation midpoint, *c*_m_, was defined as the inflection point of the individual sigmoidal curve (see **Data analysis and statistics**, below).

### Kinetic measurements

Protein was pre-incubated in the desired buffer in a 1.5 ml Protein LoBind tube (Eppendorf). Buffer that would be added to the protein was premixed in a 1.5 ml microcentrifuge tube. Refolding (or unfolding) was initiated by manually mixing the buffer with the protein and transferring the resulting mixture to the cuvette inside of the fluorometer, resulting in a 10 s dead time. Data collection started precisely 10 s after initial mixing of buffer and protein. Data was collected by excitation at 280 nm (slit widths 2 nm) and acquiring emission at 335 nm and 350 nm (slit widths 5 nm) simultaneously using two photomultiplier detectors. Corrected 335/350 was used to represent folding kinetics. An inconsistency between the kinetics measured in this work and previous work [19, 22] was determined to arise due to differences in the protein storage condition used in previous work, which did not successfully preserve the native conformation of P.69T, leading to a subpopulation of PFS in the starting sample. This heterogeneity of sample was then insufficiently unfolded at the beginning of the refolding experiments because of the extremely slow unfolding kinetics of PFS.

### Data analysis and statistics

Single exponential models were fit to the following equation:

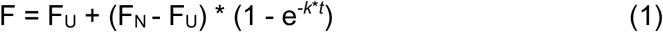

where F represents the measured fluorescence at each time point, F_U_ and F_N_ represent the fluorescence of fully unfolded protein (U) and native protein (N), *k* represents the rate constant and *t* represents time.

Double exponential models were fit to the following equation:

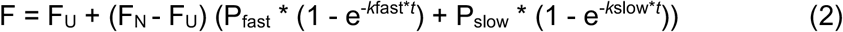

where P_fast_, P_slow_, *k*_fast_ and *k*_slow_ represent the fractional amplitude and rate constants of the fast and slow phases, respectively. Data were fit to a double exponential model only when the correlation coefficient (R^2^) of the fit to a single exponential model was lower than 0.97.

Sigmoidal models were fit to the following equation:

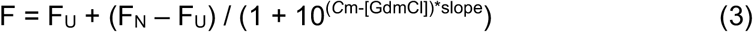

where *c*_m_ represents the GdmCl concentration at the denaturation midpoint and slope was determined by the fitting software (GraphPad Prism). For thermal denaturation experiments, *c*_m_ and [GdmCl] are replaced with melting temperature (T_m_) and temperature (T).

Double sigmoidal models were fit to the following equation:

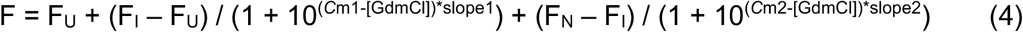

where F_I_ represents the fluorescent intensity of the baseline between the two unfolding transitions (I), *c*_m1_ and *c*_m2_ represent the GdmCl concentrations at the denaturation midpoint of transitions at lower and higher concentrations of denaturant, respectively.

Fraction native was defined by the following equation:

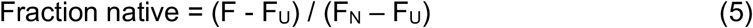

## Data Availability

Plasmids constructed for this study and raw data (spectra and kinetic traces) are available upon request from the corresponding author.

## Acknowledgements

The authors are grateful to Tobin Sosnick and members of the Clark lab for helpful discussions and Kristina Davis for assistance with creating Figure 6.

