## Supplemental Information for "Discovery of an on-pathway protein folding intermediate illuminates the kinetic competition between folding and misfolding"

Qing Luan & Patricia L. Clark*
Department of Chemistry & Biochemistry, University of Notre Dame, Notre Dame, IN 46556

### Contents

**Supplementary Tables:** Table S1 to S3

**Supplementary Figures:** Figures S1 to S8

**Table S1.** Unfolding rate constants of P.69T-Nt and P.69T-Ct, determined by fitting results shown in **Figure 3A**. For P.69T-Nt, results were fit to a single exponential equation (**Methods**, equation (1)). For P.69T-Ct, unfolding kinetics were fit to the double exponential equation (**Methods**, equation (2)); the rate constant for the slow phase is reported here.

| | **$k_U$ ($s^{-1}$)** | | | |
|---|---|---|---|---|
| **GdmCl (M)** | **P.69T-Nt** | **P.69T-Ct** | **$R^2$** | **SE from fitting** |
| 1.1 | $5.4 \times 10^{-3}$ | - | 0.9915 | $7 \times 10^{-5}$ |
| 1.3 | $1.2 \times 10^{-2}$ | - | 0.9852 | $2 \times 10^{-4}$ |
| 1.5 | $2.7 \times 10^{-2}$ | - | 0.98876 | $4 \times 10^{-4}$ |
| 1.7 | $4.3 \times 10^{-2}$ | - | 0.9783 | $8 \times 10^{-4}$ |
| 1.9 | $6.3 \times 10^{-2}$ | - | 0.874 | $3 \times 10^{-3}$ |
| 2.1 | $1.1 \times 10^{-1}$ | - | 0.8699 | $5 \times 10^{-3}$ |
| 2.3 | - | $1.2 \times 10^{-5}$ | 0.9332 | $2 \times 10^{-4}$ |
| 2.5 | - | $5.7 \times 10^{-4}$ | 0.9973 | $5 \times 10^{-6}$ |
| 2.7 | - | $2.0 \times 10^{-3}$ | 0.9975 | $2 \times 10^{-5}$ |
| 2.9 | - | $4.1 \times 10^{-3}$ | 0.9974 | $5 \times 10^{-5}$ |
| 3.1 | - | $1.6 \times 10^{-2}$ | 0.9943 | $2 \times 10^{-4}$ |

**Table S2.** Unfolding rate constants of native, PFS and PFS* P.69T and native PCt to U, determined by fitting results shown in **Figure 3B, 4C and S7**. All results were fit to a single exponential equation (**Methods**, equation (1)).

| **GdmCl (M)** | **Starting state** | **$k_U$ ($s^{-1}$)** | **$R^2$** | **SE from fitting** |
|---|---|---|---|---|
| 2.8 | Native P.69T | $2.6 \times 10^{-3}$ | 0.9963 | $2 \times 10^{-5}$ |
| 2.8 | PFS* P.69T[1] | $2.3 \times 10^{-3}$ | 0.997 | $2 \times 10^{-5}$ |
| 2.8 | PFS* P.69T-L219E[2] | $3.0 \times 10^{-3}$ | 0.9607 | $7 \times 10^{-5}$ |
| 7 | PFS P.69T | $3.0 \times 10^{-3}$ | 0.9949 | $3 \times 10^{-5}$ |
| 7 | Native PCt | $2.3 \times 10^{-3}$ | 0.9983 | $1 \times 10^{-5}$ |

[1]Native P.69T was incubated in 1.5 M GdmCl for 3 min.
[2]Fully unfolded P.69T-L219E was refolded in 0.5 M GdmCl for 10 min.

**Table S3.** Folding rate constants from U and PFS* P.69T, U P.69T-L219E and U PCt to N, determined by fitting results shown in **Figure 5B**. All results were fit to the double exponential equation (**Methods**, equation (2)); the average rate constants for the slow phase of 4~5 replicates and standard deviation were summarized in the table.

| **GdmCl (M)** | **Starting state** | **$k_U$ ($s^{-1}$)** | **Standard deviation** |
|---|---|---|---|
| 0.5 | PFS* P.69T | $5.3 \times 10^{-3}$ | $4 \times 10^{-4}$ |
| 0.5 | U P.69T | $5.8 \times 10^{-4}$ | $3 \times 10^{-5}$ |
| 0.5 | U P.69T-L219E | $7.2 \times 10^{-4}$ | $1 \times 10^{-5}$ |
| 0.5 | U PCt | $9.5 \times 10^{-4}$ | $4 \times 10^{-5}$ |

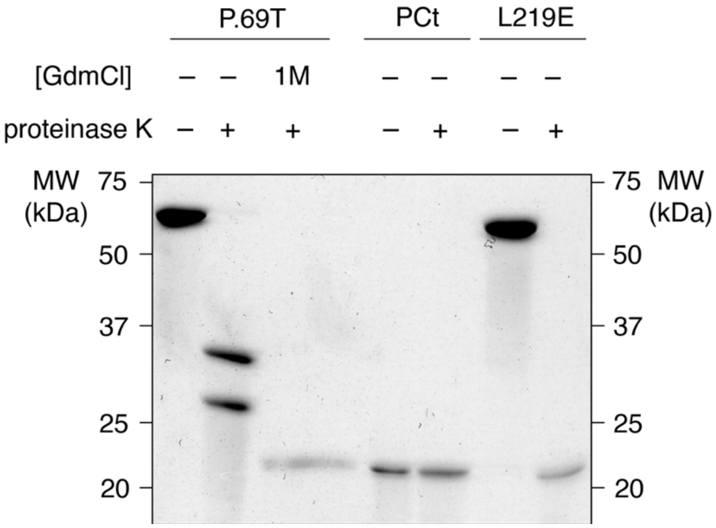


**Figure S1.** Proteinase K digestion of P.69T, PCt and P.69T-L219E. In each experiment, 5 μM protein was incubated with 2 μg/mL proteinase K for 24 hours at room temperature before SDS-PAGE and Coomassie staining. In 1 M GdmCl, P.69T adopts the PFS conformation. Native P.69T was cleaved into two fragments between 25 and 37 kDa. PFS was digested to a fragment of the same size as PCt (~21 kDa). PCt was resistant to protease digestion under these conditions.

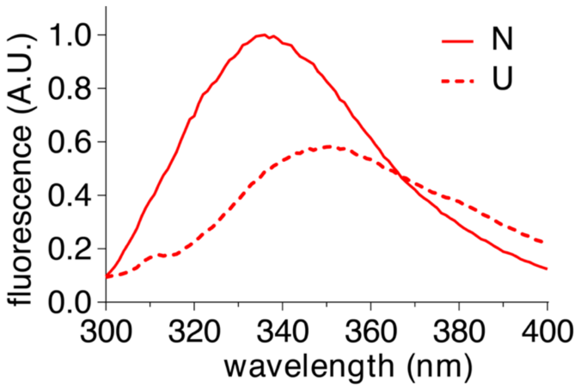


**Figure S2.** Tryptophan fluorescence emission spectra for 1 μM PCt in the native (N; 25 mM Tris HCl pH7.5, 50 mM NaCl) and chemically denatured (U; 8 M GdmCl, 25 mM Tris HCl pH7.5, 50 mM NaCl) conformations. Fluorescence emission intensity was normalized to the maximum value for the native spectrum. The small peak at 310 nm corresponds to the water Raman signal.

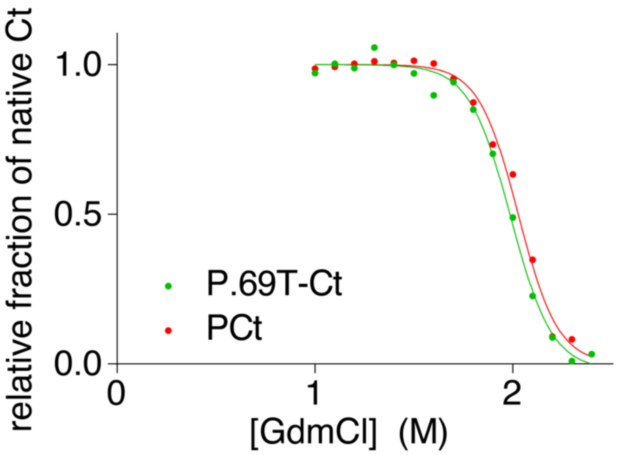


**Figure S3.** Comparison of denaturation transitions of the Ct of P.69T and PCt at equilibrium. PCt refolding at 6 days was used to represent equilibrium. The tryptophan fluorescence emission ratio between 335 and 350 nm was calculated and normalized to the ratio for native Ct. For this analysis, the pre-transition baseline for P.69T between 1-1.5 M GdmCl was defined as 100% native for P.69T-Ct.

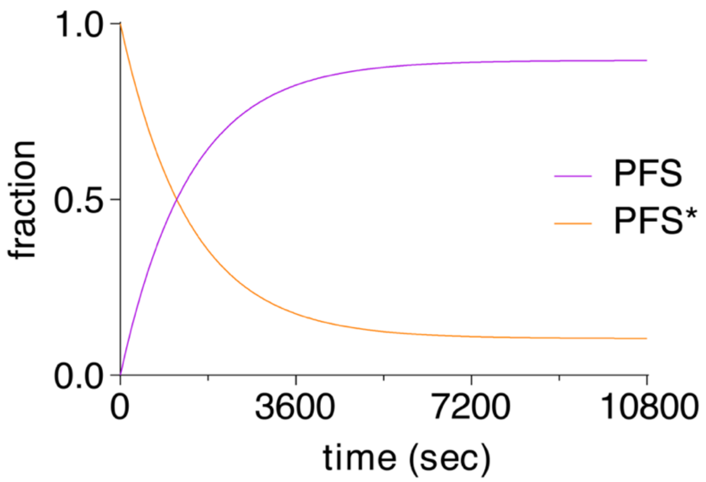


**Figure S4.** Accumulation kinetics of PFS* and PFS during Nt unfolding. The fractions of PFS* and PFS at each time point were calculated from **Fig 4A** and fit to a single exponential model. Only the fitted curves are shown here.

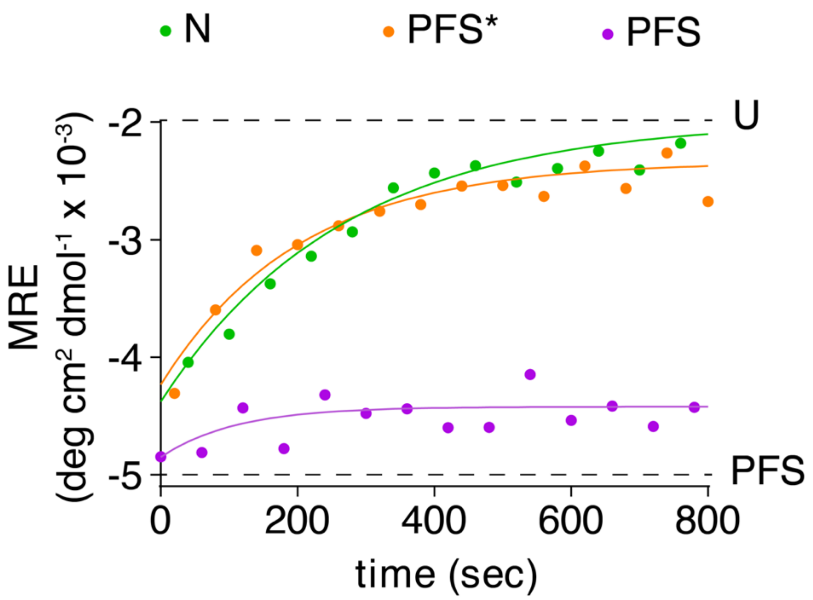


**Figure S5.** Unfolding kinetics of P.69T, PFS* and PFS in 2.8 M GdmCl monitored by far-UV CD spectroscopy at 218 nm. Native P.69T and PFS* (populated by unfolding native P.69T in 1.5 M GdmCl for 3 minutes) showed similar unfolding kinetics, whereas no unfolding was observed for PFS (populated by unfolding native P.69T in 1.5 M GdmCl overnight). The results were consistent with the kinetic experiments monitored by tryptophan fluorescence (**Figure 3B**).

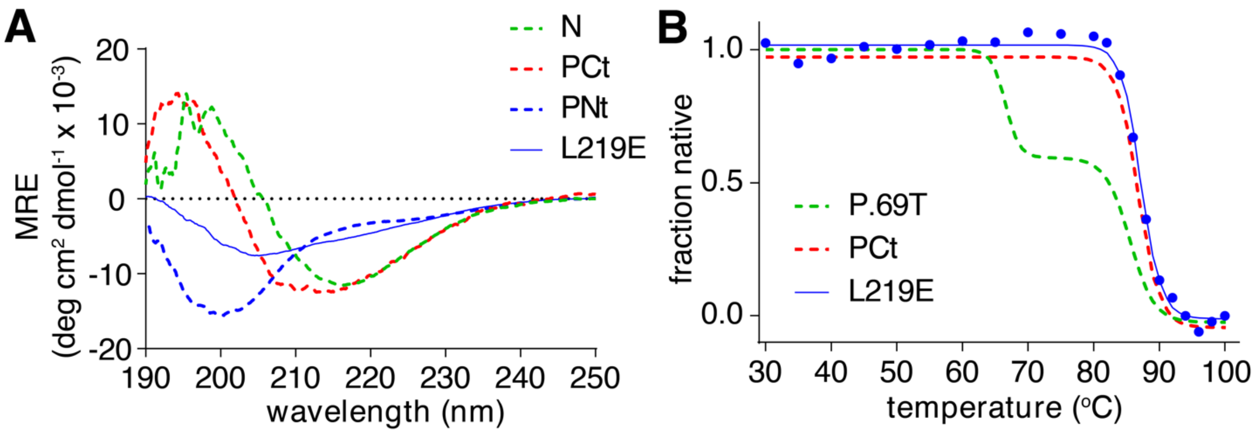


**Figure S6.** P.69T-L219E adopts a PFS-like conformation even in the absence of denaturant.
(**A**) Far-UV CD spectra of P.69T (folded; *green dashed*), PCt (folded; *red dashed*), PNt (disordered; *blue dashed*) and P.69T-L219E (*blue solid*). The spectrum of P.69T-L219E showed higher disordered content than P.69T.
(**B**) Thermal denaturation monitored by far-UV CD spectroscopy at 218 nm. The denaturation of P.69T-L219E (*blue*) was fitted to a single sigmoidal, consistent with PCt (*red*) and the second unfolding transition of P.69T (*green*) (see also **Figures 1 & S4**).

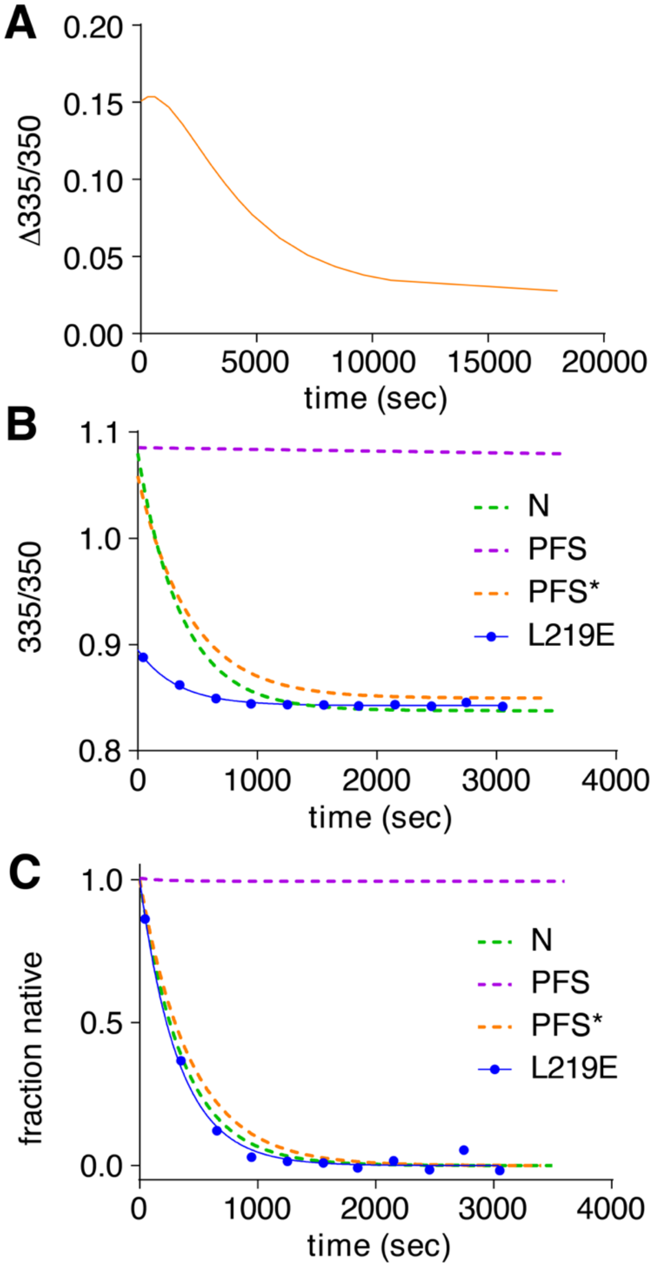


**Figure S7.** Unfolding kinetics of the PFS*-like structure populated during P.69T-L219E folding is similar to that of PFS* populated from P.69T unfolding.
(**A**) Apparent kinetics of P.69T-L219E in PFS* presented as the hatched region in **Fig 5C**.
(**B**) P.69T (*green dashed*), PFS (*purple dashed*), PFS* (*orange dashed*) and P.69T-L219E PFS* (*blue solid*) were unfolded in 2.8 M GdmCl. All curves were fit to a single exponential model. Only fitted curves were shown for the dashed lines.
(**C**) Converting the curves in (**B**) to fraction native to directly compare the kinetics.

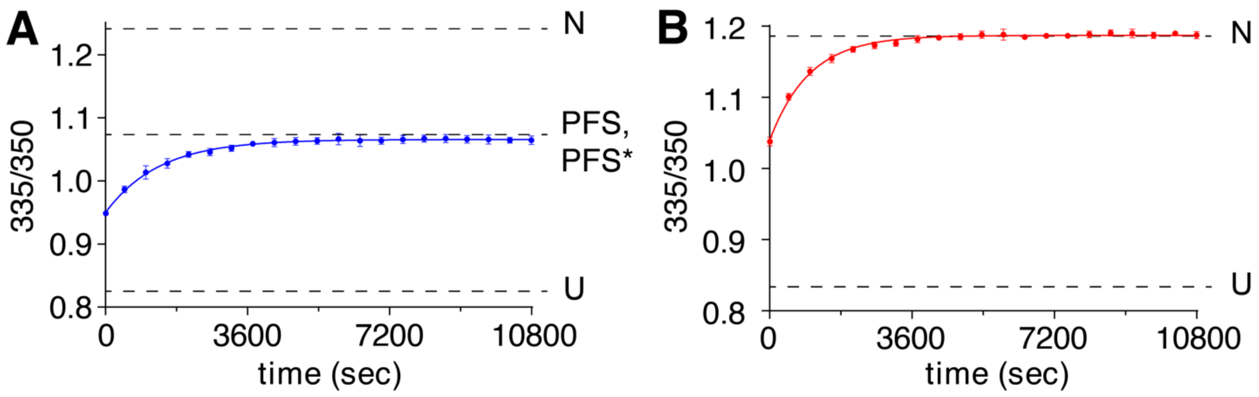


**Figure S8.** Folding kinetics of P.69T-L219E (**A**) and PCt (**B**). Protein was fully unfolded in 7 M GdmCl for 1 hour before dilution to 0.5 M GdmCl to initiate refolding. Refolding was monitored by tryptophan fluorescence emission. Data were fit to a double exponential, with a fast phase in the dead time. The folding rate constants of the slow phases is compared to the fully unfolded and PFS* conformations of wild type P.69T in **Figure 5B**.
